# A β-importin promotes fusome growth and distribution in *Drosophila* ovarian germ cells

**DOI:** 10.1101/2025.03.10.642432

**Authors:** Amanda M. Powell, Anna E. Williams, Elizabeth T. Ables

## Abstract

Germline cysts, or interconnected groups of germ cells, promote the synchronization of gamete development in vertebrates and invertebrates. In the *Drosophila* ovary, cyst formation is coordinated by the fusome, a unique endoplasmic reticulum-like organelle. Although many structural components of the fusome have been characterized, little is known about the genetic factors that control fusome growth and distribution between germ cells. Here, we identify the β-importin, *Tnpo-SR*, as an important regulator of fusome morphogenesis in germline stem cells (GSCs) and their dividing daughters. Although Tnpo-SR does not aggregate at fusomes or centrosomes, *Tnpo-SR* null mutants fail to form proper fusomes and knock-down of *Tnpo-SR* reduces fusome accumulation, ultimately leading to cyst fragmentation and improper numbers of germ cells within an egg chamber. *Tnpo-SR* depletion disrupts microtubule organization during interphase and sub-cellular localization of the microtubule associated protein Asp. However, overexpression of *asp* is not sufficient to restore fusome size or cyst architecture when *Tnpo-SR* is depleted, suggesting that *Tnpo-SR*-dependent regulation of *asp* does not solely control fusome morphogenesis. Instead, we find that restoring fusome size by overexpressing the core fusome component *hu-li tai shao* or the polarity factor *Par3/bazooka* is sufficient to restore fusome area in *Tnpo-SR*-depleted GSCs. Moreover, restoration of fusome area in the absence of *Tnpo-SR* also rescues cyst organization and oocyte specification, suggesting that *Tnpo-SR* functions upstream of fusome structural component production. Taken together, these data functionally link nuclear import/export machinery to fusome morphogenesis during *Drosophila* germline cyst development.

**ARTICLE SUMMARY:** The fusome plays a key role in cyst formation in the *Drosophila* ovarian germline, yet regulation of fusome growth and distribution between germ cells after mitosis remains understudied. Through loss-of-function analyses, the authors identify *Tnpo-SR* as a novel regulator of fusome morphogenesis. *Tnpo-SR* depletion reduces the core fusome structural component Hts in germline stem cells (GSCs) and disrupts microtubule organization and nucleation. Overexpression of *Hts* or *Par3/bazooka* in *Tnpo-SR*-depleted GSCs restores fusome size, which in turn rescues cyst organization and oocyte specification. These findings establish Tnpo-SR as an upstream regulator of Hts during fusome morphogenesis.

## INTRODUCTION

Gametogenesis is a highly conserved biological process that allows multicellular organisms to produce similar specialized sex cells (Lesch and Page 2012; Nicholls and Page 2021; Spradling et al. 2022). Female gametes, or oocytes, provide the zygote with a haploid genome and the cellular machinery required to initiate and guide early embryonic development (Arur 2017; Elkouby and Mullins 2017; Spradling et al. 2022). To ensure oocytes accumulate sufficient maternal material, many vertebrate and invertebrate species form interconnected clusters of germ cells, called germline cysts, which support efficient gamete production (de Cuevas et al., 1997; Kloc et al., 2004; Levy et al., 2024; Spradling et al., 2022). Germline cysts are most well studied in adult *Drosophila*, where each 16-cell cyst gives rise to a single oocyte (Figure 1A-B) (Hinnant et al., 2020). Following their formation, cysts are surrounded by somatic follicle cells and encapsulated into individual egg chambers (Figure 1A). Evidence from *Drosophila* and mouse suggests that cyst formation provides structural and mechanical support, promotes temporal synchronization of gamete development, and enables oocyte quality control (Lu et al. 2017; Ikami et al. 2023; Spradling 2024; Levy et al. 2024).

**Figure 1:**
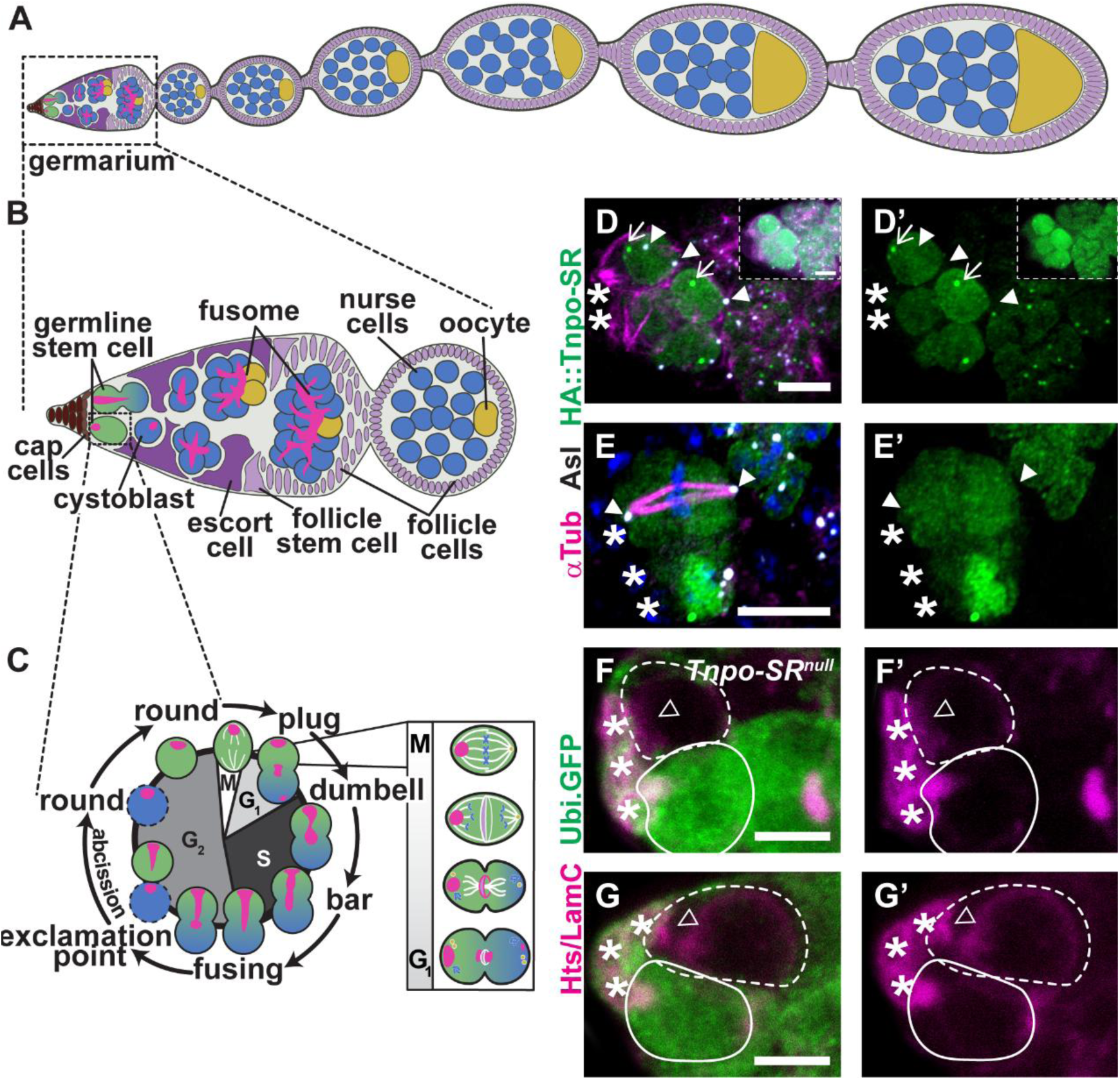
Tnpo-SR does not aggregate in fusomes or centrosomes, but null mutants of *Tnpo-SR* fail to form fusomes. (A) *Drosophila* ovariole. Progressively older egg chambers contain 16 cells [15 nurse cells (blue) and 1 oocyte (yellow)]. (B) Germaria lie at the anterior most tip of the ovariole, where somatic cap cells (brown) are juxtaposed to GSCs (green). Asymmetric division of a GSC yields a cystoblast (blue) which divides four times to produce a 16-cell cyst. The fusome (magenta) interconnects germ cells. (C) GSC fusome morphogenesis parallels progression through the cell cycle. (D-E) Maximum intensity projections of interphase (D) and mitotic (E) *UASz-HA::Tnpo-SR* GSCs immunostained for Tnpo-SR (HA, green), microtubules (⍺-Tubulin, magenta), centrosomes (Asl, white) and DNA (DAPI, blue). Insets in panels (D-D’) are the same germarium imaged with (full) and without (inset) Airyscan high resolution imaging with deconvolution. (F-G) Mosaic germaria harboring homozygous null *Tnpo-SR^KG^* GSCs (visualized by the absence of GFP, green, dotted outline) and control GSCs (visualized by the presence of GFP, green, solid outline) immunostained for Hts (fusome, magenta) and LaminC (nuclear membrane, magenta). Scale bars = 5µm.

Foundational studies in the *Drosophila* ovary also underscore the importance of a unique endoplasmic reticulum (ER)-like organelle, the fusome, in coordinating cyst cell divisions and specifying the oocyte. This germline-specific organelle is conserved across arthropods and some vertebrates, including *Xenopus* and mouse (Röper and Brown 2004; Kloc et al. 2004; Nakamura et al. 2010; Nakamura et al. 2011; Elkouby and Mullins 2017; Davidian and Spradling 2025; Pathak and Spradling 2026). Early work in *Drosophila* reports that germ cells lacking cytoskeletal *β-Spectrin* or the adducin homolog *hu li tai shao* (*hts*) do not form functional fusomes (as visualized by ⍺-Tubulin and ⍺-Spectrin) and cysts fail to divide past the 4-cell stage (Lin et al. 1994; de Cuevas et al. 1997; Deng and Lin 1997). Similarly, loss of the microtubule associated protein Orbit or the recycling endosome protein Rab11 induces defects in fusome elongation and morphogenesis, ultimately disrupting cyst architecture (Bogard et al. 2007; Miyauchi et al. 2013). Moreover, the fusome organizes the polarized microtubule network necessary to specify the oocyte. Loss of minus-end associated *Patronin* (*CAMSAP/ssp4*), the plus-end associated *mini spindles* (*msps/XMAP215*), or the actin crosslinker *short stop* (*shot*) leads to cysts with 16 polyploid germ cells and lacking an oocyte (Theurkauf et al. 1993; Lu et al. 2021; Nashchekin et al. 2021; Lu et al. 2023; Nashchekin et al. 2024; Barr et al. 2024; Lu et al. 2026). Together, these data support the model that the fusome coordinates synchronous cyst divisions by acting as a trafficking hub and microtubule organizer. Interestingly, not all fusome-associated proteins are required for organelle expansion or for maintaining cyst synchronicity, suggesting a hierarchy of essential versus dispensable components (Lighthouse et al. 2008). Although *Drosophila* provides the most detailed understanding of fusome composition and spatiotemporal dynamics, the molecular mechanisms that guide its growth and distribution during cell division remain unclear.

In the *Drosophila* ovary, cyst fusomes originate in the germline stem cell (GSC), where the organelle is assembled, segregated, and re-established at each cell cycle (Figure 1B-C) (Hinnant et al., 2020; Telfer, 1975). Early electron micrographs show that fusome material accumulates as mitotic spindle remnants condense at the intracellular bridge (Telfer 1975). More recent fluorescence imaging confirms that ⍺-Tubulin persists at the fusome throughout interphase even after mitotic spindle remnants have disassembled, likely at lower concentrations or in a monomeric state (Grieder et al. 2000; Williams and Ables 2023). For most of the cell cycle, GSCs contain a large, round fusome (sometimes called the spectrosome) anchored at the anterior margin near neighboring somatic cap cells (de Cuevas et al. 1997; Huynh 2006; Ong et al. 2010; Ables and Drummond-Barbosa 2013; Villa-Fombuena et al. 2021). During asymmetric division, a small plug of fusome material is deposited into the differentiating daughter cell (the prospective cystoblast; also termed the pre-cystoblast) (de Cuevas et al. 1997; Huynh 2006; Ong et al. 2010; Ables and Drummond-Barbosa 2013; Villa-Fombuena et al. 2021). The fusome then expands between the GSC and cystoblast as they remain interconnected through G_2_ of the next cell cycle (Figure 1C) (Ables and Drummond-Barbosa 2013; Villa-Fombuena et al. 2021). After abscission in late G_2_, both the GSC and cystoblast fusomes revert to a round morphology in preparation for the next division. The cystoblast subsequently undergoes four incomplete mitotic divisions to form a 16-cell cyst. During GSC and cyst divisions, the fusome anchors one centrosome per cell to maintain synchrony while extending through all cells to form its characteristic branched structure. In early cysts (2- and 4-cell), fusomal ER is continuous with cytoplasmic ER (Snapp et al. 2004). As the cyst matures, ER continuity is lost, and the fusome is largely disassembled. However, the polarized microtubule array established by the fusome remains essential for transporting maternal determinants into the oocyte even after organelle dissolution (Starz-Gaiano and Lehmann 2001; Bastock and St Johnston 2008; Hinnant et al. 2020).

We previously characterized the β-importin *Tnpo-SR* as an essential regulator of female germline cyst development (Beachum et al. 2023). In the absence of *Tnpo-SR*, fusome size is reduced in GSCs and cysts, and fusome continuity is impaired. Here, we use *Tnpo-SR*-depleted germ cells to investigate how fusomes are partitioned from GSCs into the subsequent germ cell lineage, and how a β-importin, an essential regulator of cell division, contributes to this process. We find that *Tnpo-SR*, like other β-importins, is required to exclude the microtubule-binding protein Abnormal Spindle (Asp) from the nucleus and to inhibit microtubule growth during interphase. However, *asp* overexpression does not restore fusome size or cyst defects in *Tnpo-SR*-depleted germ cells. Instead, our data suggest that *Tnpo-SR* promotes fusome distribution by indirectly impacting accumulation of the core fusome component Hts. Overexpressing *hts* in *Tnpo-SR*-depleted germ cells restores fusome area and significantly improves cyst formation and oocyte specification. These data reveal new connections between the mitotic cell cycle machinery and fusome morphogenesis and clearly demonstrate how alterations to fusome architecture impact cyst formation and subsequent oocyte specification. More broadly, our studies add to the growing body of evidence that β-importins perform specialized, context-dependent roles in coordinating intracellular organization.

## MATERIALS AND METHODS

### *Drosophila* Strains and Husbandry

Flies were maintained at 22-25°C in standard medium (cornmeal/molasses/ yeast/agar) (NutriFly MF; Genesee Scientific). A complete list of fly strains used are listed in the Key Resources Table. Female progeny were collected within 24 hours of eclosion and maintained on wet yeast for three days prior to dissection at 25°C, unless otherwise indicated.

*Tnpo-SR* null GSCs (Figure 1F-G) were generated by inducing genetic mosaics using *Flippase (FLP)/FLP recognition target (FRT)* mitotic recombination, as previously described (Beachum et al. 2023). For RNAi experiments, germline knockdown of *Tnpo-SR* was facilitated by expressing the *pValium22*-based transgene *UAS-Tnpo-SR^RNAi^* using the germline-specific *nos-GAL4::VP16-nos.UTR* (Doren et al. 1998; Beachum et al. 2023). Females carrying the driver and one UAS element were used as controls.

*Tnpo-SR^KI^* is a loss-of-function allele resulting from CRISPR-mediated insertion of *RFP-loxP-3xP3-GFP-alpha-Tub-3’UTR-loxP* into the open reading frame of the *Tnpo-SR* locus (Figure S1) (Well Genetics, Inc.). In contrast to our previously published *mCherry::Tnpo-SR* (Beachum et al. 2023), *Tnpo-SR^KI^* flies are homozygous lethal and do not express RFP.

### Immunofluorescence and Microscopy

Ovaries were prepared for immunofluorescence microscopy as described (Hinnant et al. 2017; Williams and Ables 2023). Ovaries were dissected and teased apart in room temperature Grace’s medium without additives (Caisson Labs) and fixed in 5.3% formaldehyde in Grace’s medium for 10 min at room temperature. They were then washed extensively in phosphate-buffered saline (PBS, pH 7.4; Fisher) with 0.1% Triton X-100, permeabilized in PBS with 0.5% Triton X-100 for 30 minutes and blocked for 1-3 hours in blocking solution [5% normal goat serum (MP Biomedicals), and 0.1% Triton X-100 in PBS] at room temperature. Primary antibodies (listed in the Key Resources Table) were diluted in blocking solution and used overnight at 4°C. Secondary antibodies were followed by a 2-hour (or overnight) incubation at room temperature with AlexaFluor 488- 568- or 633-conjugated goat species-specific antibodies (Life Technologies; 1:200). All ovary samples were stained with 0.5 μg/ml DAPI (Sigma) in 0.1% Triton X-100 in PBS and mounted in 90% glycerol mixed with 20% n-propyl gallate (Sigma).

Confocal z-stacks (0.5-1.0 μm optical sections) were collected with the Zeiss LSM700 laser scanning microscope or Zeiss LSM800 laser scanning microscope with Airyscan detector (Zen software). Airyscan images were subject to deconvolution protocols (Zen). Images were analyzed using Zen or FIJI and minimally and equally enhanced via histogram using ZEN and Adobe Photoshop Creative Suite.

### Statistical Analysis

GSCs were identified based on the juxtaposition of their fusomes to the junction of adjacent cap cells. Fusome measurements were taken from the center-most z-stack (or maximum projection if the fusome was split between z-stacks) of each GSC. The average area of the fusome was collected using the FIJI area tool, measuring the pixel area of the identified fusome. Asymmetric plug measurements were taken from each piece of the plug (mother/daughter) and plotted as paired data points. GSC loss was measured as the average number of GSCs per germaria.

Mitotic spindle measurements were taken using the line tool in FIJI using a max z-stack projection of the mitotic GSC being analyzed (from the bottom of the cell to the top). Length measurements were collected from one centrosome to the other (identified by centrosome marker Asl). Width measurements were taken from the center of the mitotic spindle at its widest point.

Interphase microtubules were identified based on the presence of ⍺-Tubulin emanating from centrosomes (identified by centrosome marker Asl). A Fisher’s Exact test was performed using Prism (Graphpad) comparing driver controls versus *driver>UAS-RNAi*.

For analyses involving the egg chambers outside of the germarium, data was collected from egg chambers posteriorly located from the germarium. Oocytes were identified the presence of anti-Orb antibodies and a condensed nuclei. The number of cells were the number of cell nuclei visualized with DAPI. Results were subjected to Student’s two-tailed T-test comparing driver controls versus *driver>UAS-asp, hts, baz* or *driver>UAS-RNAi* versus *driver>UAS-RNAi*, *UAS-asp*, *hts* or *baz*.

All results were evaluated using Microsoft excel and Prism (GraphPad), subjected to either a Student’s two-tailed T-test or Fisher’s exact test, where \*\*\**p*<0.00001, \*\**p*<0.0001, \**p*<0.001. (In Figure S2, a lower statistical threshold was set to accommodate lower sample sizes; here, \**p*<0.01).

## RESULTS

### *Tnpo-SR* promotes proper fusome morphogenesis in GSCs

We previously showed that germ cells depleted of *Tnpo-SR* formed egg chambers with irregular cell numbers and produced cysts with small, thin fusomes (Beachum et al. 2023). Because loss of other fusome-associated factors, such as *hts* and *α-Spectrin*, also results in abnormal cyst cell numbers (Lin et al. 1994; de Cuevas et al. 1996), we hypothesized that these phenotypes might be functionally linked and that Tnpo-SR may localize to the fusome to coordinate this function. We analyzed intracellular localization using high-resolution microscopy and a hemagglutinin-tagged *Tnpo-SR* transgene expressed in germ cells (Figure 1D-E). Consistent with other β-importins, Tnpo-SR localized to both the cytoplasm and nucleus during interphase, with higher concentrations in the nucleus and in discrete puncta (Figure 1D-D’). Notably, Tnpo-SR did not co-localize with fusomes or with the centrosomal protein Asterless (Asl) (Figure 1D-E). During mitosis, Tnpo-SR was distributed throughout the cell, consistent with diffusion through the semi-permeable nuclear envelope, and did not co-localize with DNA, microtubules, or spindle centrosomes (Figure 1E-E’).

Given the localization of Tnpo-SR, we were surprised to find that GSCs homozygous for a null mutation of *Tnpo-SR* were almost completely devoid of fusomes, as visualized with Hts (Figure 1F-G). This observation, along with the analysis of Tnpo-SR localization, suggested that Tnpo-SR might indirectly regulate fusome morphogenesis with subsequent effects on fusome deposition in cyst daughters. Unfortunately, *Tnpo-SR* null GSCs were often found as single cells, indicating they eventually stopped dividing (Beachum et al. 2023). We therefore turned to two independent hypomorphic genetic models: females carrying germline-specific short hairpin RNA (*Tnpo-SR^RNAi^*) and females heterozygous for a mild loss-of-function insertion (*Tnpo-SR^KI^*; Figure S1A). Importantly, *Tnpo-SR^RNAi^* females produced egg chambers with abnormal germ-cell numbers, whereas *Tnpo-SR^KI^* heterozygotes produced largely normal egg chambers (Figure S1B) (Beachum et al. 2023). Using these models, we measured Hts-positive fusome area in GSCs at comparable stages, grouping fusomes into “round” (G_2_/M) and “stretched” (G_1_/S and early G_2_) categories (Figure 2). Unlike *Tnpo-SR* nulls, fusomes from all stages were present in both hypomorphs, and *Tnpo-SR^RNAi^* GSCs progressed through the cell cycle largely normally, albeit with a longer G_1_ phase (Figure S2). At eclosion, depleting *Tnpo-SR* via RNAi or the knock-in allele showed an average **∼**0.7-fold reduction in fusome area compared with age-matched wild-type GSCs in the same fusome class (Round=0.632, Stretched=0.762; *p*<0.00001) (Figure 2A-H).

**Figure 2:**
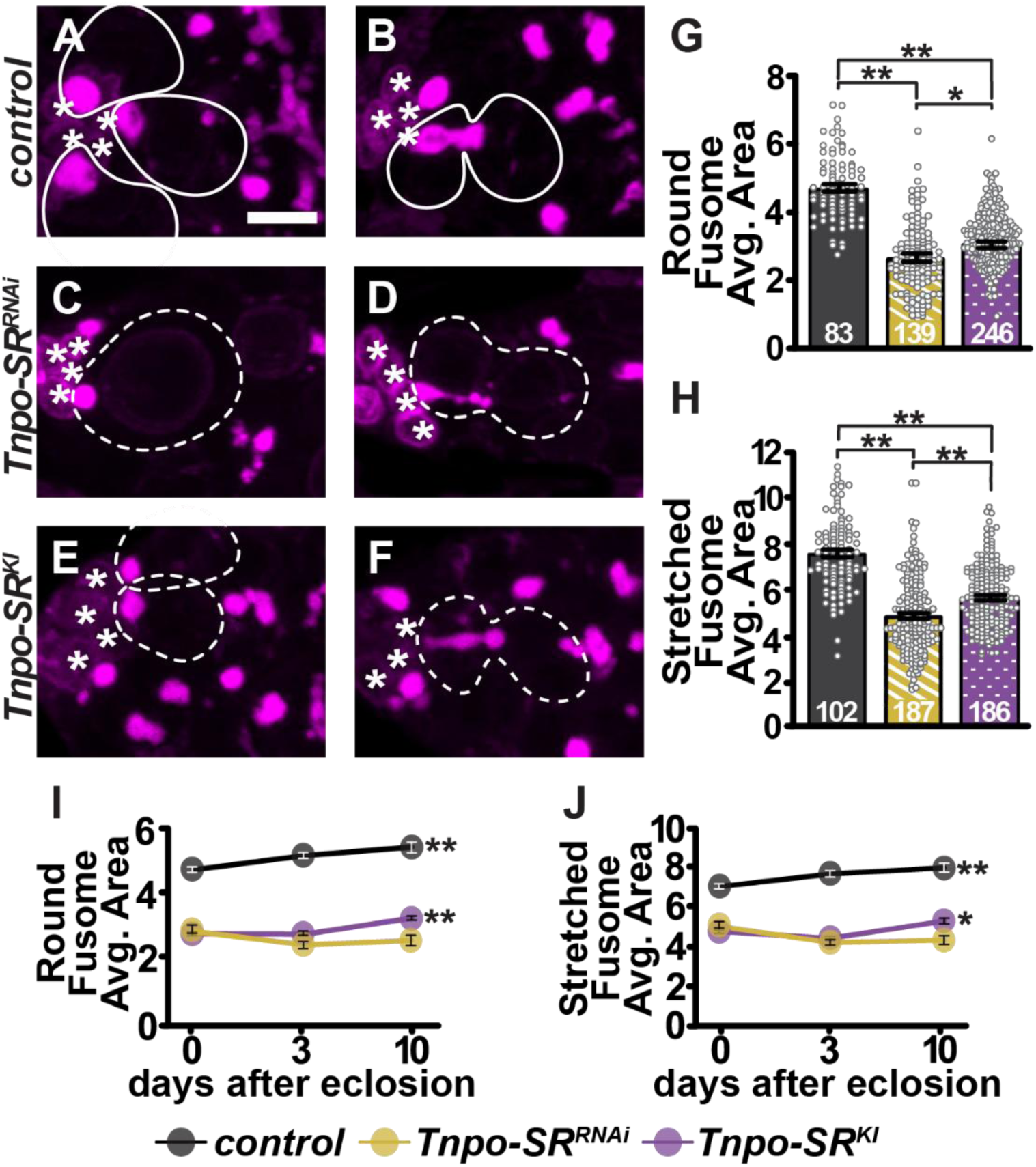
*Tnpo-SR* promotes fusome growth. (A-F) Max projections of germaria from control (A-B), *Tnpo-SR^RNAi^* (C-D) and *TnpoSR^KI^* (E-F) ovaries immunostained for the fusome (Hts, with LamC to mark cap cells, magenta). Asterisks denote cap cells; GSCs outlined with white lines (solid–control, dashed–mutant). (G-H) Quantification of the fusome area in round (G) and stretched (H) fusome morphologies at 6 days after eclosion. (I-J) Quantification of fusome area in round (I) and stretched (J) fusome morphologies over time. Significance determined by Student’s two-tailed T-test between genotypes (G-H) or between ages (0dae vs. 10dae) of the same genotype (I-J). Numbers in bars represent the number of GSCs analyzed. Error bars represent s.e.m. \*\**p*<0.0001, \**p*<0.001. Scale bar = 5µm.

To further characterize the hypomorphic models, we asked whether fusome size changed over successive cell divisions. Intriguingly, wild-type GSC fusomes grew larger from eclosion to ten days after eclosion, while *Tnpo-SR^RNAi^* GSC fusomes did not (Figure 2I-J). Surprisingly, *Tnpo-SR^KI^* GSC fusomes, although smaller at eclosion, increased slightly over time, more closely resembling the wild-type trajectory (Figure 2I-J). These data suggest that *Tnpo-SR* supports fusome growth as females age.

We then asked whether other fusome-associated proteins had a similarly reduced area at the fusome in *Tnpo-SR*-depleted GSCs. We first visualized another core fusome component, ⍺-Spectrin, and saw a similar reduction in fusome size compared to Hts at six days after eclosion (Figure 3A-B, E-F). We next quantified two membrane-associated proteins, Reticulon-like1 (Rtnl1) and the apicobasal protein Scribble (Scrib) and again observed comparable reduction between control and *Tnpo-SR*-depleted GSCs (Figure 3). Since Hts is necessary for proper accumulation of ⍺-Spectrin and Rtnl1 (Yue and Spradling 1992; Lin et al. 1994; de Cuevas et al. 1996; Röper 2007; Lighthouse et al. 2008), we conclude that reduction in Hts impacts the levels of other fusome-associated proteins. Together, these data indicate that *Tnpo-SR* is essential for establishing proper fusome size in GSCs. Moreover, because fusomes were reduced in size but not absent, *Tnpo-SR* hypomorphs provide a novel and useful tool for dissecting fusome morphogenesis.

**Figure 3:**
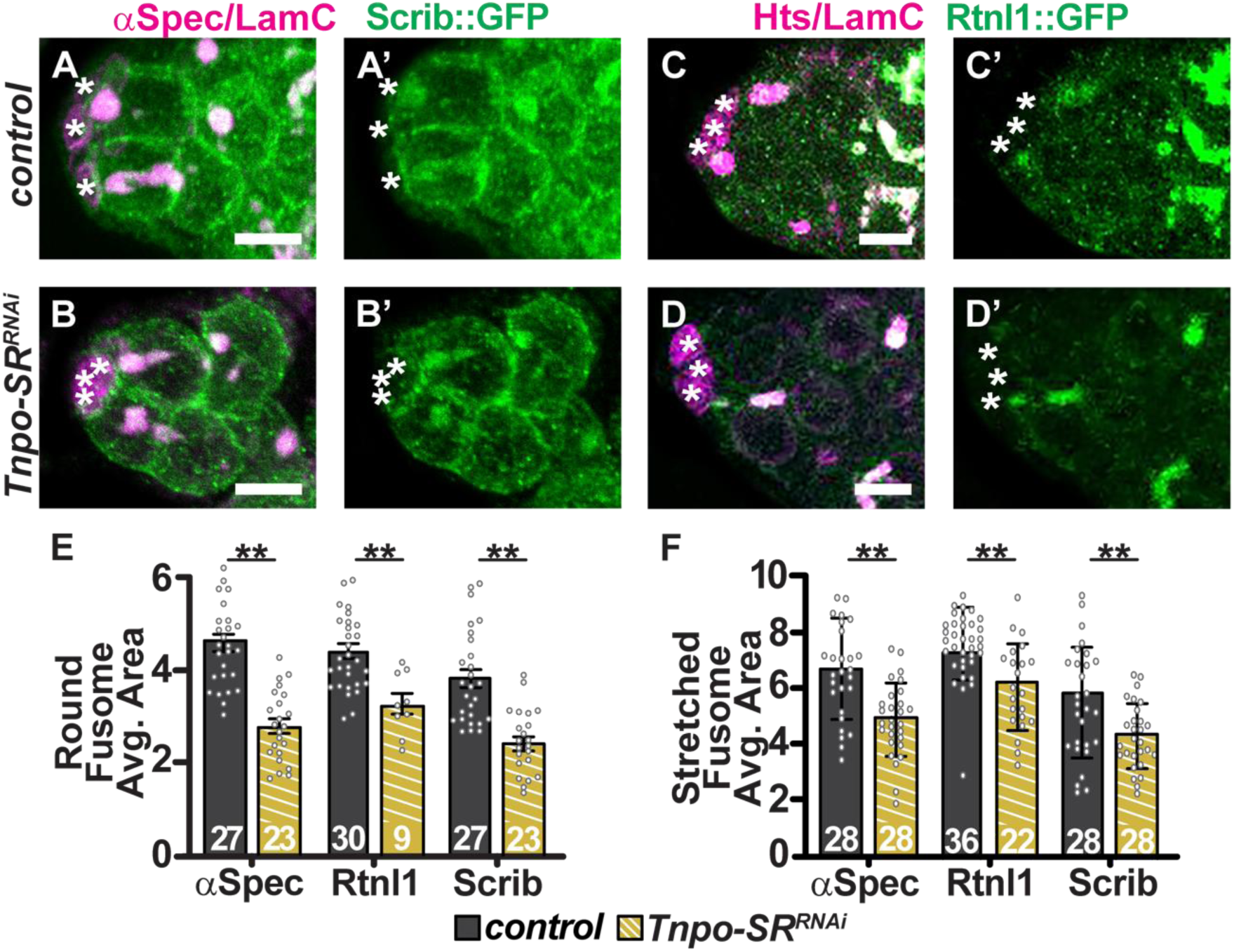
Reduction of Hts in *Tnpo-SR*-depleted GSCs is accompanied by reduction in other fusome core and fusome-associated proteins. (A-D) Max projections of germaria from *Scrib::GFP* (green, A-B) or *Rtnl1::GFP* (green, C-D) control females (A,C) or recombined with *Tnpo-SR^RNAi^* (B, D), co-stained for fusome core proteins ⍺-Spectrin (A-B, magenta) or Hts (C-D, magenta) (with LamC with visualize cap cells). Asterisks (*) denote cap cells. Scale bar = 5µm. (E-F) Quantification of fusome area in round (E) and stretched (F) fusome morphologies using different markers of the fusome. Numbers in bars represent the number of fusomes analyzed. Error bars represent s.e.m. Significance determined by Student’s two-tailed T-test \*\**p*<0.0001.

Through careful quantification of Hts area and comparison with the well-described fusome morphogenetic cycle, we observed an unusual distribution of Hts in *Tnpo-SR*-depleted GSCs. In wild-type GSCs, fusome material accumulated at the cytokinetic furrow, forming the characteristic “plug” morphology that denotes the beginning of G_1_ (Figure 4A). At this stage, fusome distribution was uneven, as more Hts accumulated at the apical pole than the posterior pole, which sits at the constriction point between the GSC and presumptive cystoblast. When *Tnpo-SR* was reduced, Hts distribution was disrupted; the daughter (plug) fusome became larger than the apical (mother) fusome (Figure 4C,E). Plotting the two fusome parts as paired data points showed that wild-type pairs produced negatively sloped lines, whereas *Tnpo-SR*-depletion produced a marked increase in cystoblast-biased distribution (positively sloped lines, magenta) (Figure 4G). Despite the fusome distribution defect at G_1_, the fusome was still largely GSC-biased throughout the remainder of the cell cycle, and *Tnpo-SR*-depleted cystoblasts still expanded the fusome across divisions, reminiscent of branching in wild-type cysts (Diegmiller et al. 2023). We speculate that the imbalance or misdirection in fusome growth contributes to the weak, thin, and disorganized fusomes that are characteristic of *Tnpo-SR* null cysts (Beachum et al. 2023).

**Figure 4:**
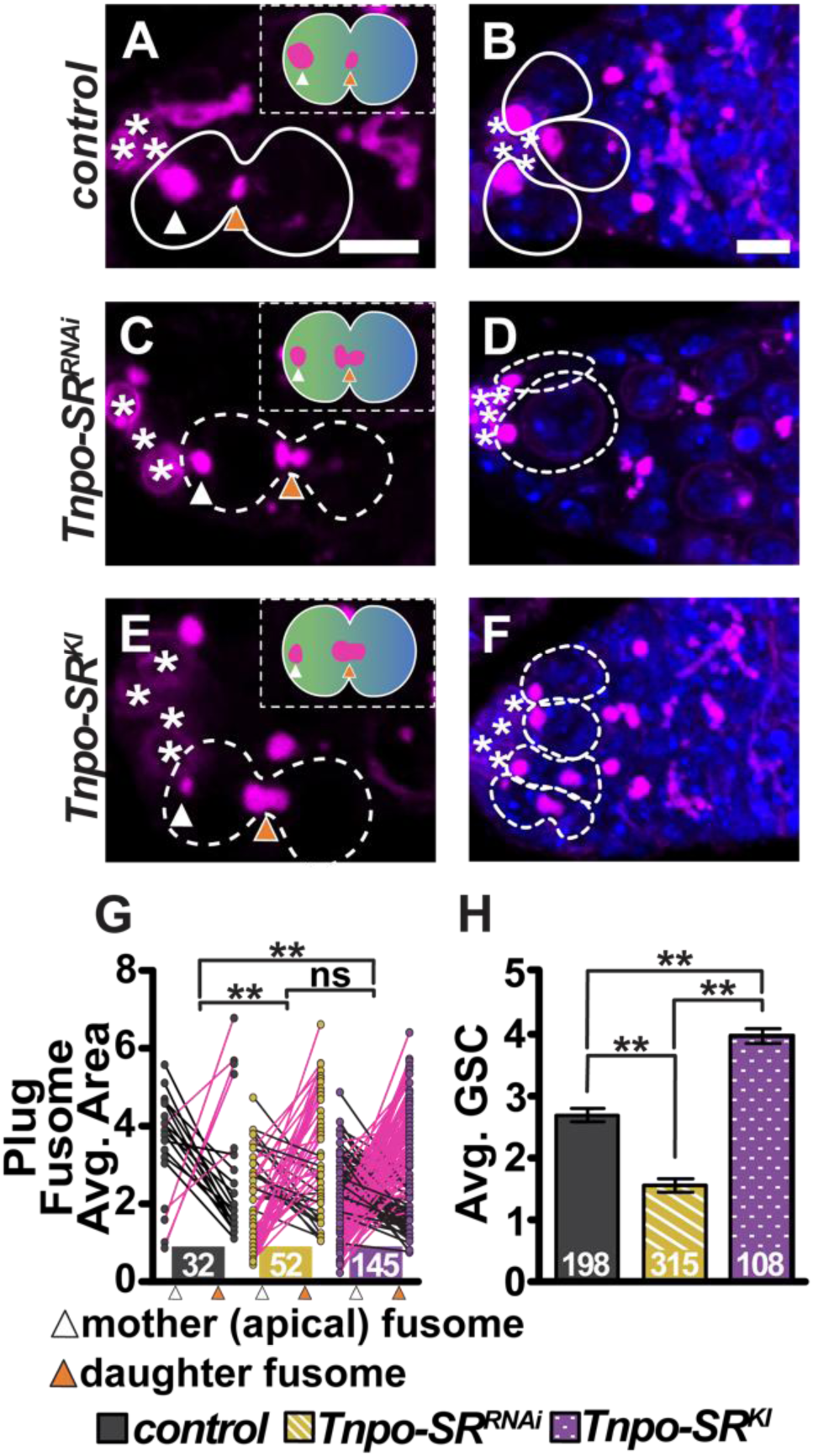
*Tnpo-SR* is required for proper fusome distribution and GSC population size. (A-F) Max projections of germaria from control (A-B), *Tnpo-SR^RNAi^* (C-D), and *Tnpo-SR^KI^* (E-F) ovaries immunostained for the fusome (Hts, with LamC to mark cap cells, magenta) and DNA (DAPI, blue). Asterisks denote cap cells; GSCs outlined with white lines (solid–control, dashed–mutant). (G) Quantification of ‘mother’ (white) and ‘daughter’ (orange) fusome size in a plug morphology. Magenta lines represent GSCs where the daughter fusome was larger than the mother fusome (positively sloped). (H) Quantification of the average number of GSCs per germaria. Numbers in bars represent the number of GSCs (G) or germaria (H) analyzed. Error bars represent s.e.m. Significance determined by Fisher’s Exact test (G) or Student’s two-tailed T-test (H); \*\**p*<0.0001. Scale bar = 5µm.

We previously described *Tnpo-SR* as a necessary regulator of GSC establishment and maintenance (Beachum et al. 2023). We wondered whether the fusome itself was important for GSC self-renewal. In fusome-less mutants, this question is difficult to address because the fusome is also a key marker used to identify GSCs. Our newly characterized *Tnpo-SR^KI^* hypomorph provided insight into this unresolved question. *Tnpo-SR* null mutants and RNAi-mediated knockdown yielded similar phenotypes in which GSCs fail to be maintained in the niche over time (Figure 4B,D,H) (Beachum et al. 2023). However, in heterozygous *Tnpo-SR^KI^* mutants (which have smaller fusomes, similar to *Tnpo-SR^RNAi^*, see Figure 2G-H), we observed a significant increase in the average number of GSCs at six days after eclosion (Figure 4D,H). Taken together, these findings indicate that *Tnpo-SR* is necessary for proper fusome morphogenesis, distribution, and integrity, but that reduced fusome accumulation alone is insufficient to compromise maintenance of the stem cell pool.

### Depletion of *Tnpo-SR* causes aberrant interphase centrosomal microtubules

β-importins have been attributed a variety of roles during mitosis, including regulation of mitotic spindle assembly, at least in part through association and sequestration of microtubule associated proteins (Nachury et al. 2001; Wiese et al. 2001; Askjaer et al. 2002; Chen et al. 2015; Beaudet et al. 2017; Beaudet et al. 2020). Since fusome regeneration in GSCs begins during the final stages of mitosis, as new fusome material forms from the remnants of the mitotic spindle (Lin et al. 1994; Villa-Fombuena et al. 2021), we hypothesized that *Tnpo-SR* could impact mitotic spindle assembly or disassembly and thus fusome morphogenesis. However, visualization of the mitotic spindle using α-Tubulin and analysis of spindle function revealed no significant differences between control and *Tnpo-SR*-depleted GSCs. Spindle length and width during mitosis were equivalent (Figure S3A-B,I-J), and accumulation of the spindle stabilizing matrix proteins Chromator and Megator was similar between control and *Tnpo-SR*-depleted GSCs (Figure S3E-H). *Tnpo-SR*-depleted GSCs still formed central spindles at the end of mitosis (Figure S3C-D), and there were no significant differences in central spindle width, suggesting that disassembly of the spindle was not impaired by reduction of *Tnpo-SR*. Finally, quantification of centromere number (visualized with CID/CENP-A antibodies) revealed no significant differences between *Tnpo-SR*-depleted and control GSCs, suggesting normal chromosome segregation (Figure S4A-C). Together, these data suggest that the role of *Tnpo-SR* in fusome growth is independent of putative activity in mitotic spindle assembly or function.

Since importins have also been shown to inhibit microtubule growth during interphase (Nachury et al. 2001; Wiese et al. 2001; Cavazza and Vernos 2016), we next focused our analysis on microtubule organization outside of mitosis. We visualized microtubules and the core fusome protein Hts in control and *Tnpo-SR*-depleted GSCs expressing fluorescently-tagged Scrib (Scrib::GFP), which is enriched at the cell membrane and in the fusome in GSCs (Lighthouse et al. 2008). Because the GSC/cystoblast pair undergoes delayed cytokinesis and remains connected via a temporary intracellular bridge where fusome proteins accumulate, Scrib::GFP allowed us to clearly visualize microtubule distribution between the GSC and cystoblast (Figure 5). In control GSCs, ⍺-Tubulin localized to the fusome and cell cortex throughout interphase and GSC fusomes changed shape in parallel to cell cycle progression as previously described (Figure 5A-G) (Villa-Fombuena et al. 2021). In contrast, *Tnpo-SR*-depleted GSCs accumulated disorganized microtubule bundles as early as G_1_ (Figure 5H-N). Intracellular interphase ⍺-Tubulin fibers failed to colocalize with the fusome or cell membranes, and occasionally oriented away from the fusome (see Figure 5N-N’, as example).

**Figure 5:**
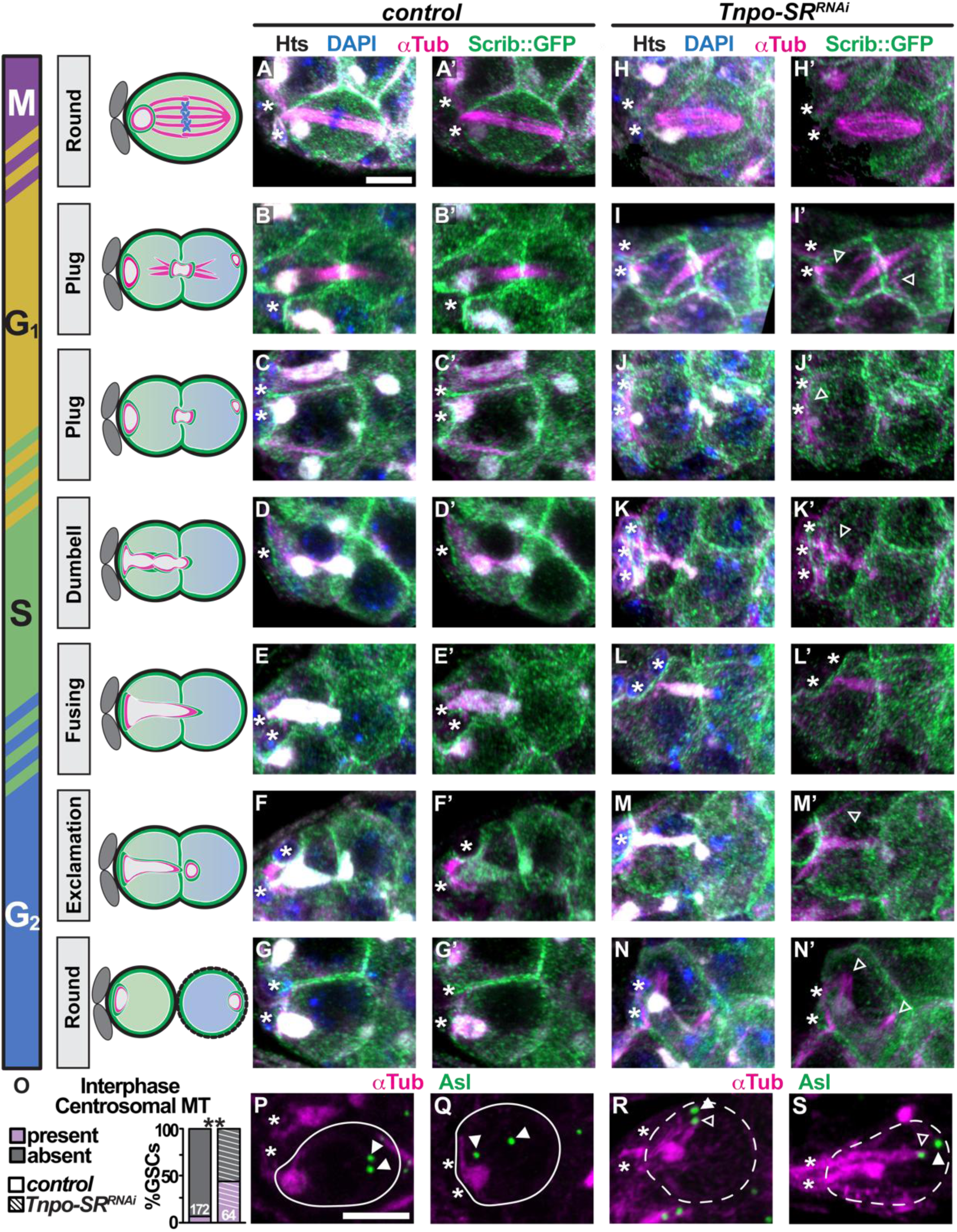
*Tnpo-SR* maintains microtubule organization during interphase. Control (A-G, P-Q) and *Tnpo-SR^RNAi^* (H-N, R-S) GSCs immunostained for the fusome (Hts, white), microtubules (⍺-Tubulin, magenta), the cell membrane (Scrib::GFP, green) or centrosomes (Asl, green, P-S), and DNA (DAPI, blue). Graphics represent the wild-type fusome morphological changes and the subsequent localization of Hts, Scrib and ⍺-Tubulin as GSCs go through the cell cycle. Asterisks denote cap cells, GSCs outlined with white lines (solid-control, dashed-mutant). Open arrowheads denote microtubule defects (M’, N’, R, S), solid arrowheads denote centrosomes (P-S). (O) Quantification of interphase GSCs where microtubules emanate from centrosomes. Significance determined by Fisher’s Exact Test, \*\**p*<0.0001. Scale bar = 5µm.

These observations suggested that *Tnpo-SR* may suppress microtubule growth from the centrosomes during interphase. To test this, we co-localized α-Tubulin and the centrosome protein Asl (Figure 5O-S). In *Tnpo-SR*-depleted GSCs, microtubules emanating from centrosomes were located throughout the cytoplasm, oriented toward the somatic cap cells, and occasionally extended protrusions that wrapped around cap cells (Figure 5O, R-S). These findings support a role for *Tnpo-SR* in suppressing centrosome-driven microtubule growth during interphase. Interestingly, despite the strong association of centrosomes with the fusome in dividing cysts (Deng & Lin 1997, Yamashita & Fuller 2008), we did not observe consistent fusome-centrosome association in control or *Tnpo-SR*-depleted GSCs, indicating that *Tnpo-SR* function in suppressing microtubule growth is independent of the molecular mechanisms that position centrosomes in preparation for mitotic spindle assembly (Figure S4C-H). Taken together, these results suggest that improper microtubule organization and growth accompany the reduction in fusome size observed in *Tnpo-SR*-depleted GSCs.

### Sub-cellular localization of Asp during interphase requires *Tnpo-SR*

Given its putative role as a nucleocytoplasmic transporter and the microtubule-disorganization phenotype associated with loss of *Tnpo-SR*, we hypothesized that *Tnpo-SR* might control the stability, transport, or translation of a protein essential for microtubule dynamics in germ cells. We first screened a variety of microtubule associated proteins by visualizing their localization in *Tnpo-SR^RNAi^* germ cells. Among these, we observed consistent mis-localization of Abnormal Spindle (Asp), a microtubule-binding protein that localizes to the poles of mitotic spindles and the minus ends of microtubules. Previous studies demonstrated that *asp* is essential for proper germ cell divisions and oocyte specification and functions as a regulator of microtubule nucleation (Saunders et al. 1997; Do Carmo Avides and Glover 1999; Riparbelli et al. 2004). Unfortunately, we were unable to source anti-Asp antibodies to visualize endogenous Asp. Instead, we overexpressed a GFP-tagged version of *asp* specifically in germ cells and confirmed its localization at the poles and centrosomes of mitotic spindles (Figure S5A). At the telophase-to-G_1_ transition, Asp was excluded from the reforming nuclear membrane and remained localized at the poles of the central spindle, but was notably absent from the nucleus and the plug of fusome material at the constriction point (Figure S5B-C). During the rest of interphase, Asp was clearly excluded from the nucleus and enriched in the cytoplasm and fusome (Figure 6A).

**Figure 6:**
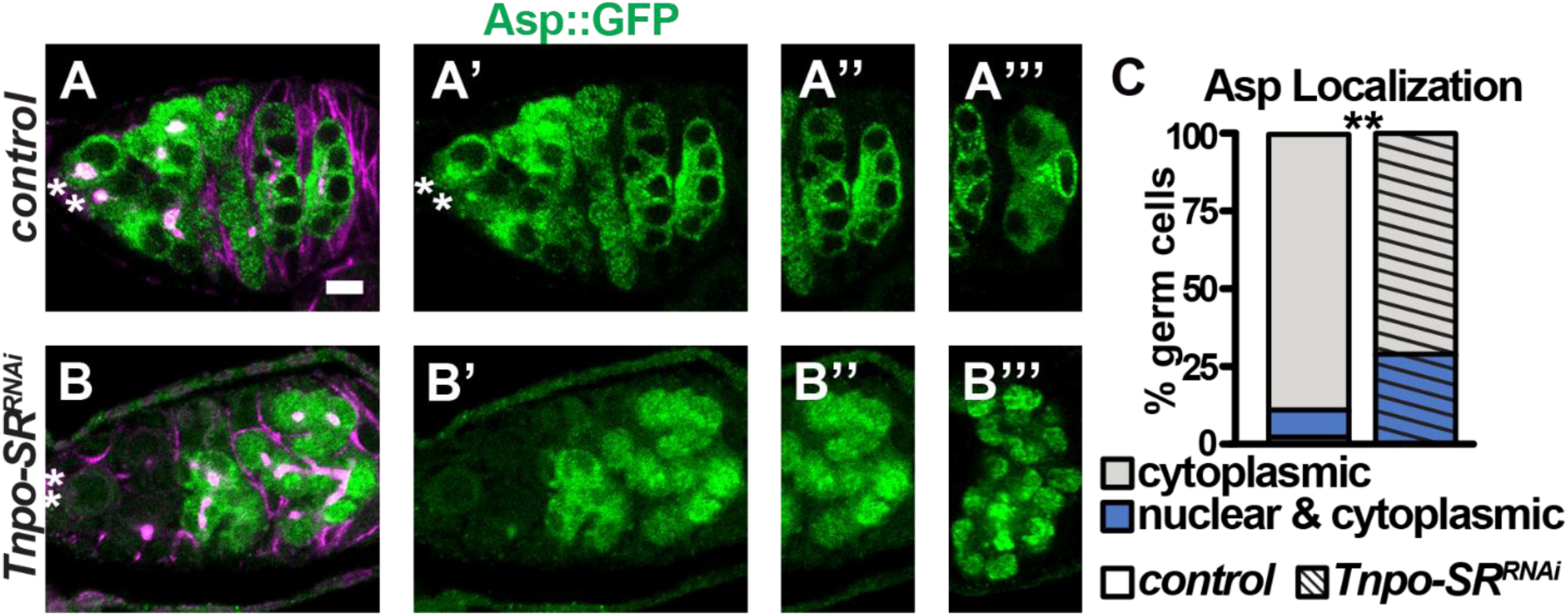
*Tnpo-SR* directs the sub-cellular localization of Asp during interphase. (A-B) *UASp-Asp::GFP* (A-A’’’) and *Tnpo-SR^RNAi^* (B-B’’’) germaria immunostained for Asp (GFP, green) and the fusome (Hts, with LamC to mark cap cells, magenta). (C) Quantification of cysts with Asp localized to the cytoplasm (gray) or to the nucleus and cytoplasm (blue). Asterisks mark cap cells. Significance determined by Fisher’s Exact Test, \*\**p*<0.0001. Scale bar = 5µm.

In mitotic germ cells depleted of *Tnpo-SR*, Asp clearly localized at the centrosome and poles of mitotic spindles. However, during interphase, we observed a significant number of germ cells where Asp localized to both the cytoplasm and nucleus (Figure 6B-C). Although we occasionally observed wild-type 4- and 8-cell cysts with cytoplasmic and nuclear Asp localization (Figure 6C), we speculate that these cysts are in the early stages of mitosis when the nuclear membrane is semi-permeable. Since *Tnpo-SR*-depleted germaria do not have a significant increase in the number of mitotic cysts, it is unlikely that the increased number of *Tnpo-SR*-depleted germ cells with abnormal nuclear Asp localization is due to alterations in the cell cycle. Further, we never observed nuclear localization of Asp in wild-type 16-cell cysts (that have exited the mitotic cell cycle), whereas nuclear Asp localization could be readily identified in 16-cell cysts depleted of *Tnpo-SR* (compare Figure 6A’’’ and 6B’’’). These results suggest that *Tnpo-SR* is necessary to retain or stabilize Asp in the cytoplasm during interphase.

### Restoration of Hts in *Tnpo-SR*-depleted cells promotes proper cyst formation

Our *Tnpo-SR*-depletion model gave us opportunity to better understand the relationship between the establishment of the fusome in early divisions and its connection to germ cyst development. We first reasoned that if microtubules were the primary regulators of fusome morphogenesis, and if this process relies on *Tnpo-SR*, then overexpression of *asp* (a microtubule nucleator) in the absence of *Tnpo-SR* should restore GSC fusome size and subsequent germline cyst organization. Whereas depletion of *Tnpo-SR* reduced fusome size, altered the number of germ cells per egg chamber, and altered the number of oocytes per egg chamber, overexpression of *asp* alone did not significantly increase fusome size or alter the number of germ cells in an egg chamber, as compared to wild-type controls (Figure 7A,C,G-J). Intriguingly, overexpression of *asp* in *Tnpo-SR*-depleted germ cells did not increase fusome size or rescue the number of germ cells per egg chamber (Figure 7D, G-I). However, overexpression of *asp* in *Tnpo-SR*-depleted germ cells significantly improved oocyte specification (Figure 7D,J). We speculate that overexpression of *asp* may be sufficient to bias microtubule localization to a single germ cell, improving oocyte selection in the absence of *Tnpo-SR*. These data suggest that although *Tnpo-SR* promotes proper intracellular localization of Asp, this function is independent of the role of *Tnpo-SR* in fusome morphogenesis and subsequent cyst formation.

**Figure 7:**
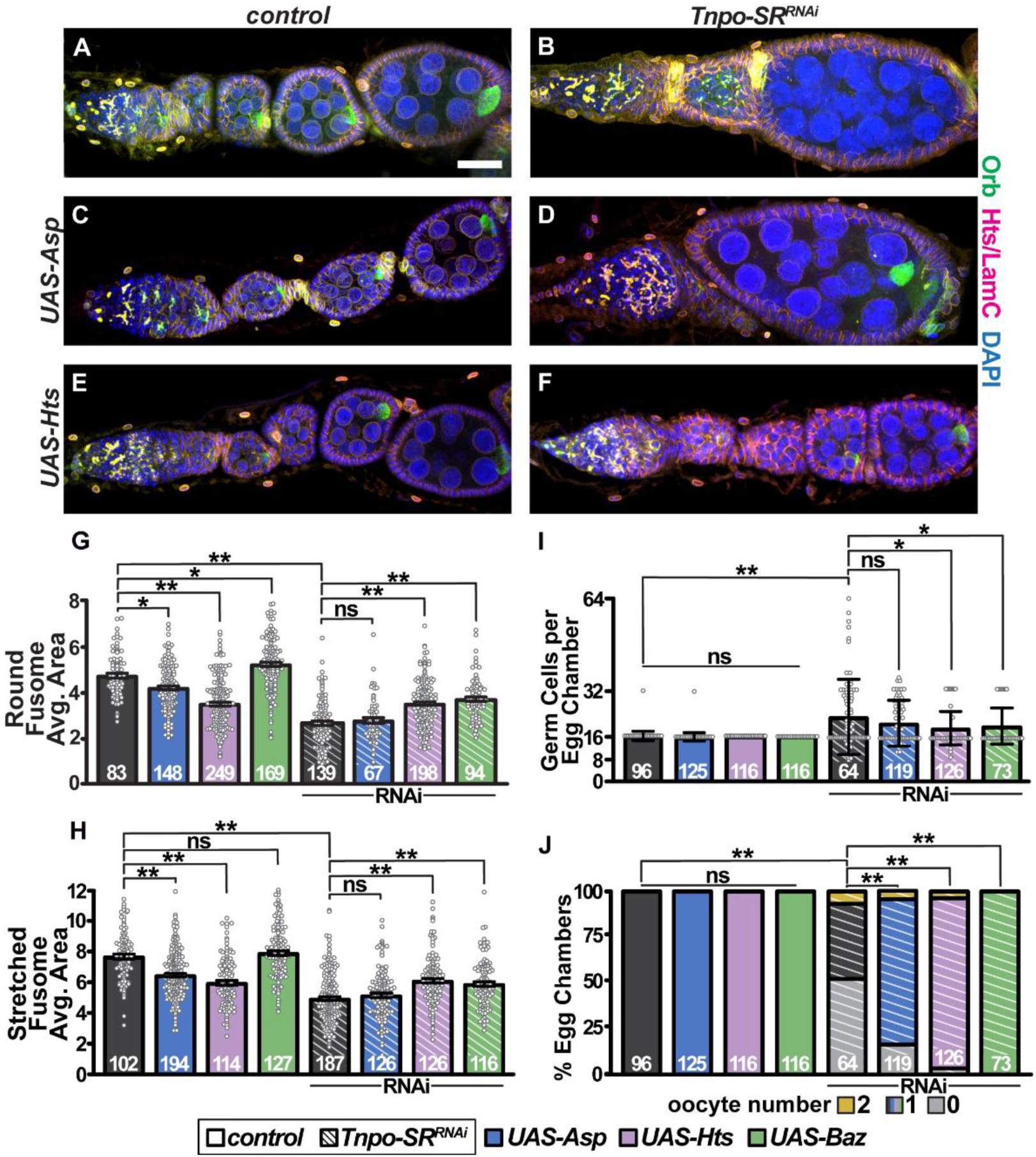
Restoration of fusome area is sufficient to rescue cyst formation in *Tnpo-SR*-depleted germ cells. (A-F) Representative ovarioles from control (A, C, E) and *Tnpo-SR^RNAi^* (B, D, F) females carrying *UASp-Asp::GFP* (C, D) or *UASp-Hts::mCherry* (E, F) immunostained for the fusome (Hts, with LamC to mark cap cells, magenta), oocytes (Orb, green), and DNA (DAPI, blue). Scale bar = 20µm. (G-H) Quantification of fusome area in round (G) and stretched (H) fusome morphologies. (I-J) Number of germ cells (I) or oocytes (J) per egg chamber. Numbers in bars represent number of GSCs (G,H) or egg chambers (I,J) analyzed. Bars indicate control (gray); *UAS-Asp* (blue); *UAS-Hts* (purple); *UAS-Baz* (green); *Tnpo-SR^RNAi^* (striped). Significance determined by Student’s two tailed T-test (G, H, I) or Fisher’s exact test (J), \*\**p*<0.0001, \**p*<0.001.

We then asked whether increasing fusome size was sufficient to restore cyst formation in *Tnpo-SR*-depleted germ cells. We discovered that overexpression of the apical polarity protein *Par3/bazooka* (*baz*) significantly increased the size of the GSC fusome in both round and stretched morphologies (Figure 7G-H). Baz localizes primarily to the apical side of GSCs, bordering the membrane between the GSC and the cap cells, and minimally at the fusome. The overexpression of *baz* alone did not impact the number of cells within an egg chamber or oocyte specification (Figure 7I-J). However, overexpression of *baz* in *Tnpo-SR*-depleted germ cells significantly increased fusome size, as compared to depletion of *Tnpo-SR* alone (Figure 7G-H). Moreover, overexpression of *baz* also significantly rescued the number of germ cells per egg chamber and oocyte specification in the absence of *Tnpo-SR*, with 81% of egg chambers containing 16 cells with one oocyte (Figure 7I-J).

Given these results, we then asked whether overexpression of *Hts* itself was sufficient to restore fusome morphogenesis in *Tnpo-SR*-depleted germ cells. Overexpression of a mCherry-tagged *hts* yielded slightly smaller fusomes than wild-type, probably because unlike endogenous Hts, a large amount of Hts-mCherry is shuttled to the cell cortex (Figure 7E, G-H). This did not impact the production of 16-cell egg chambers with a single oocyte (Figure 7E, I-J). When we overexpressed *hts* in *Tnpo-SR*-depleted germ cells, we observed a significant increase in fusome area as compared to the RNAi alone (Figure 7F, G-H). Overexpression of *hts* also significantly improved the average number of germ cells in an egg chamber, as 80% of egg chambers had the appropriate cyst composition (Figure 7F, I-J). Of the remaining egg chamber, 16% had 32 cells with two oocytes, suggesting that either two cysts were encapsulated together or that a 16-cell cyst went through fifth round of division. Together, these results demonstrate that defects in germ cyst formation caused by the depletion of *Tnpo-SR* arise primarily from impaired fusome accumulation, and that *hts* likely lies downstream of *Tnpo-SR* function in germ cells.

## DISCUSSION

The β-importin encoded by *Tnpo-SR* is essential for oogenesis in *Drosophila*. Previous studies demonstrated that *Tnpo-SR* promotes GSC self-renewal, GSC and germ cell mitotic division, cyst formation, and oocyte selection (Beachum et al. 2023). Whether these phenotypes were interconnected, or indicative of the more general role of importins as necessary regulators of mitosis, was unclear. Here, we demonstrate that *Tnpo-SR* regulates fusome morphogenesis independently of its role in cell division. We observe abnormal accumulation and distribution of the core fusome component Hts during interphase in *Tnpo-SR* hypomorphic models and link this phenotype with the establishment of proper germline cyst organization. Although we do not detect defects in mitotic spindle assembly, our data suggest that *Tnpo-SR* controls microtubule nucleation and organization during interphase, likely by limiting nuclear localization of the microtubule-associated protein Asp. Since Tnpo-SR does not accumulate at fusomes, centrosomes, or microtubules, we posit that Tnpo-SR may also regulate fusome morphogenesis indirectly by controlling localization of a splicing factor that impacts Hts protein accumulation. In support of this model, we show that restoring Hts accumulation, either directly by overexpressing *hts* or indirectly by overexpressing *Par3/baz*, rescues cyst organization and oocyte specification in the absence of *Tnpo-SR*. These findings position *Tnpo-SR* as a novel regulator of fusome morphogenesis in the *Drosophila* germline and suggest that fusome structure depends on proper nuclear-cytoplasmic compartmentalization, linking nuclear import machinery to cytoplasmic organelle biogenesis in an unexpected and previously unrecognized way. This insight may help explain how specialized organelles, such as the fusome, evolved to meet the coordination demands of germline cysts.

### The fusome as a regulator of *Drosophila* female germ cyst development

Maintaining coordination among interconnected cells must require specialized mechanisms to prevent developmental asynchrony and cyst fragmentation. We propose that the fusome evolved in *Drosophila* in concert with germline cyst development to serve this coordinating function. The architecture of the fusome reflects its dual roles: ER-like membranous vesicles facilitate protein and organelle transport between cyst cells, while the robust adducin-based cytoskeletal framework physically anchors centrosomes during divisions, ensuring mitotic synchronicity. Although a fusome-like structure has not been found in all organisms, morphologically similar organelles have been identified. In Tardigrade germ cysts, researchers noted an enormous ER-like structure that occupied the center of nurse cells (Suzuki 2006). *Xenopus* forms 16-cell cysts with a branched structure traversing the ring canals, closely resembling that of *Drosophila* cysts (Kloc et al. 2004). Most recently, fusome-like structures have also been identified in mouse germline cysts (Pathak and Spradling 2026). Despite morphological similarities between these ER-associated organelles, the presence of distinct protein compositions in each system suggests that the molecular mechanisms guiding fusome morphogenesis have diverged or evolved separately between species to accomplish a conserved function (Davidian and Spradling 2025).

Prior studies suggested that the fusome plays a central role in oocyte selection (reviewed in Hinnant et al. 2020). Yet recent experiments have shed important caveats to this model. Barr et al. (2024) found that while both pro-oocytes contain two-fold the amount of fusome material as their daughter cells, there is no association between which pro-oocyte has the most fusome material and the accumulation of the oocyte differentiation factor Orb (Barr et al. 2024). However, the physical anchoring of Orb mRNA to the fusome is a critical and necessary step in oocyte specification, as this prevents oocyte determining factors from ‘leaking’ into other cells, ensuring only one cell is specified. Although the fusome is largely disassembled by the time that egg chambers form, the specification of the oocyte, and the microtubule trafficking highway established by the fusome, persists throughout oogenesis and is necessary for the trafficking of maternal determinants between nurse cells and the oocyte (Grieder et al. 2000).

In *Drosophila* males, the fusome follows similar morphological changes as females during germline cyst division, but persists throughout spermatogenesis and is not required to maintain germline cyst development (Kaufman et al. 2020). There are a few explanations for the discrepancies in the necessity of the fusome between sexes. First, female germ cysts rely on the specification of one cell, while each cell in a male germline cyst develops into a competent spermatid. Second, female fusomes are present only during the mitotically dividing stages of oogenesis, while male fusomes persist throughout spermatogenesis. Perhaps the function of the fusome in establishing the polarized microtubule array for oocyte specification is fulfilled during mitosis, while in males the fusome acts perdures as a site for continued protein production, modification, or turnover.

### Fusome size is not required to maintain GSCs in the niche

A key unresolved question in germline biology is whether the fusome itself is essential for GSC self-renewal. Loss of function mutations that abolish the fusome become depleted of GSCs; however, this analysis is complicated by the fact that the fusome also serves as a marker of GSC identity. Our study identified a mild hypomorphic mutation of *Tnpo-SR* (*Tnpo-SR^KI^*) with reduced fusome area that phenocopies depletion of *Tnpo-SR* via RNAi and complete loss-of-function alleles. Surprisingly, *Tnpo-SR^KI^* also had more GSCs per germarium than is found in wild-type genetic backgrounds. Although additional biochemical studies are necessary, we believe these observations help us better understand the interdependent relationship between the fusome and GSC self-renewal. The fact that fusome size and GSC population size can be phenotypically separated suggests that fusome morphogenesis and GSC self-renewal are likewise separable events. This is perhaps not surprising, since GSCs rely on external signaling from the somatic cap cells to promote the activity of the bone morphogenetic protein signaling pathway and GSC self-renewal, largely independently of fusome function (Hinnant et al., 2020). Since GSCs are anchored on one side to the cap cells, they are also intrinsically and structurally biased within the tissue toward the source of BMP ligands (Song et al. 2002). BMP signaling is also necessary to maintain GSC adhesion to cap cells, thus establishing a feedback loop promoting retention of GSCs in the niche and GSC apicobasal polarity (Song et al. 2002; Song et al. 2004). Since overexpression of the apical polarity determinant *Par-3/baz* or restoring Hts expression rescued fusome area and germ cyst organization, we speculate that biased fusome distribution arises as a downstream consequence of GSC polarity. Thus, while it remains theoretically possible that localized translation of proteins within the fusome may contribute to GSC self-renewal, our study suggests that fusome size itself is not a primary driver of GSC self-renewal. Although the molecular mechanisms are separable, the fusome continues to be a useful readout of cell cycle dynamics, microtubule organization, cell polarity, and GSC identity.

### Importins function beyond nuclear transport

Importins (or transportins) have been broadly characterized as molecular protein shuttles between the nucleus and cytoplasm, taking advantage of the RanGTP gradient to power their movement (Kalab et al. 1999; Kalab et al. 2002; Mühlhäusser and Kutay 2007; Lonhienne et al. 2009; Cesario and McKim 2011). Importins are thought to directly interact with two different categories of proteins. “Cargoes” are proteins that are destined for transport to/from the nucleus, whereas “binding partners” can be transient interactions (Kuersten et al. 2001; Yang et al. 2023). Although cargoes were thought to be the main driver of importin function during interphase, more recent studies suggest that importin binding partners are necessary regulators of cellular organization during all phases of the cell cycle. In *C. elegans* and *Xenopus* embryos, Importin-β maintains the mitotic spindle through associations with Ran, although a direct binding partner was not initially identified (Wiese et al. 2001; Askjaer et al. 2002). Further studies identified the microtubule-associated protein NuMa as an intriguing binding partner of Importin-β (Nachury et al. 2001; Wiese et al. 2001). During interphase, Importin-β sequesters NuMa in the cytoplasm, suppressing ectopic microtubule assembly (Wiese et al. 2001). At the onset of mitosis, NuMa is released and trafficked to the nucleus, ensuring spindle assembly proteins are accessible near chromatin to build the mitotic spindle (Wiese et al. 2001). Further studies have termed Importin-β as a cytoplasmic chaperone, not only sequestering proteins, but also preventing abnormal protein aggregation or breakdown (Jäkel et al. 2002; Damizia et al. 2022).

The structure of Importin-β is thought to be the key to its versatility, with 19 HEAT repeats forming a spring-like superhelical structure with an inherently large degree of flexibility, allowing each interacting protein to have a unique binding site (Harel and Forbes 2004). Collectively, these studies demonstrate that Importin-β regulates cytoplasmic architecture as well as nuclear transport, providing a conceptual framework for our findings on Tnpo-SR. Like Importin-β, Tnpo-SR has 20 HEAT repeats, but to date, has only been directly implicated in the binding of phosphorylated serine/arginine (SR)-rich protein sequences, many of which are splicing factors (Allemand et al., 2002; Lai et al., 2001). Intriguingly, conformational studies on the human ortholog of Tnpo-SR, Tnpo3, have identified over one hundred potential binding partners, ranging from transcription factors to microtubule- and cell-cycle regulators (Kimura et al. 2017). More recent studies show that Tnpo3, despite the name of its *Drosophila* ortholog, is not reliant on the SR/RS repeats to define its cargoes (Zhou et al. 2025). Instead, Tnpo3 can bind a noncanonical RSY nuclear localization sequence. Using this motif, Zhou et al (2025) identified 33 additional putative cargoes, including ⍺-Adducin, the ER-associated protein Sec61⍺, and β-Spectrin.

Our analysis of *Tnpo-SR*-depleted germ cells suggests that Tnpo-SR shares functional properties with Importin-β in competitively binding microtubule associated proteins. While *Tnpo-SR* does not appear to influence mitotic spindle assembly or maintenance, it does play a significant role in the suppression of microtubules during interphase, perhaps through a direct interaction with Asp. Although Asp does not show clear SR/RS/SRY-rich protein sequences, these domains may not be necessary for transient interactions. Moreover, we show that depletion of *Tnpo-SR* directly impacts the accumulation of the adducin-like core fusome component Hts and based on Tnpo3 motif data, Tnpo-SR could directly bind Hts (Zhou et al. 2025). However, because neither interaction has been confirmed directly via protein-binding studies, and we do not see clear co-localization of Tnpo-SR at the fusome or microtubules, we cannot rule out the possibility that Tnpo-SR is a necessary transporter of a splicing factor that regulates either Asp or Hts. Further, since Hts has 16 isoforms, one of which is required for the formation of ring canals (Hts-RC), Tnpo-SR may promote localization of the splicing factors required to splice Hts, particularly if only a subset of isoforms require importin- dependent nuclear processing. Taken together, our findings support a model in which importins act as multifunctional regulatory hubs, whose cargo selection is tuned to a specific cell type and developmental context. This specialization highlights how distinct importins, despite sharing conserved interactions with the Ran GTPase system, can be selectively leveraged across cell types to coordinate cytoskeletal function and developmental programs.

## DATA AVAILABILITY STATEMENT

Strains, plasmids, and images are available upon request. The authors affirm that all data necessary for confirming the conclusions of the article are present within the article, figures, and tables.

## ACKNOWLEDGMENTS

Many thanks to the ECU Department of Biology Microscopy Core Facilities and the ECU School of Dental Medicine for microscopy support, and to members of the Ables and Anllo laboratories and our anonymous reviewers for helpful discussion and critical reading of this manuscript. *Drosophila* lines were obtained from the Bloomington *Drosophila* Stock Center (NIH P40OD018537) or Kyoto *Drosophila* stock center or kind gifts from V. Gefland, A. Spradling, D. Drummond-Barbosa, T. Harris, and B. Riggs. Antibodies were a kind gift from N. Rusan or obtained from the Developmental Studies Hybridoma Bank (DSHB), created by the NICHD of the NIH and maintained at the University of Iowa, Department of Biology.

## FUNDING

This work was supported by National Institutes of Health R15-GM117502 and R35-GM158065 (E. T. A.). A.M.P. was supported by the East Carolina University Department of Biology, Harriot College of Arts and Sciences, and Graduate School. A.E.W. was supported by the ECU Office of Undergraduate Research.

## COMPETING INTERESTS

No competing interests declared.

## KEY RESOURCES TABLE

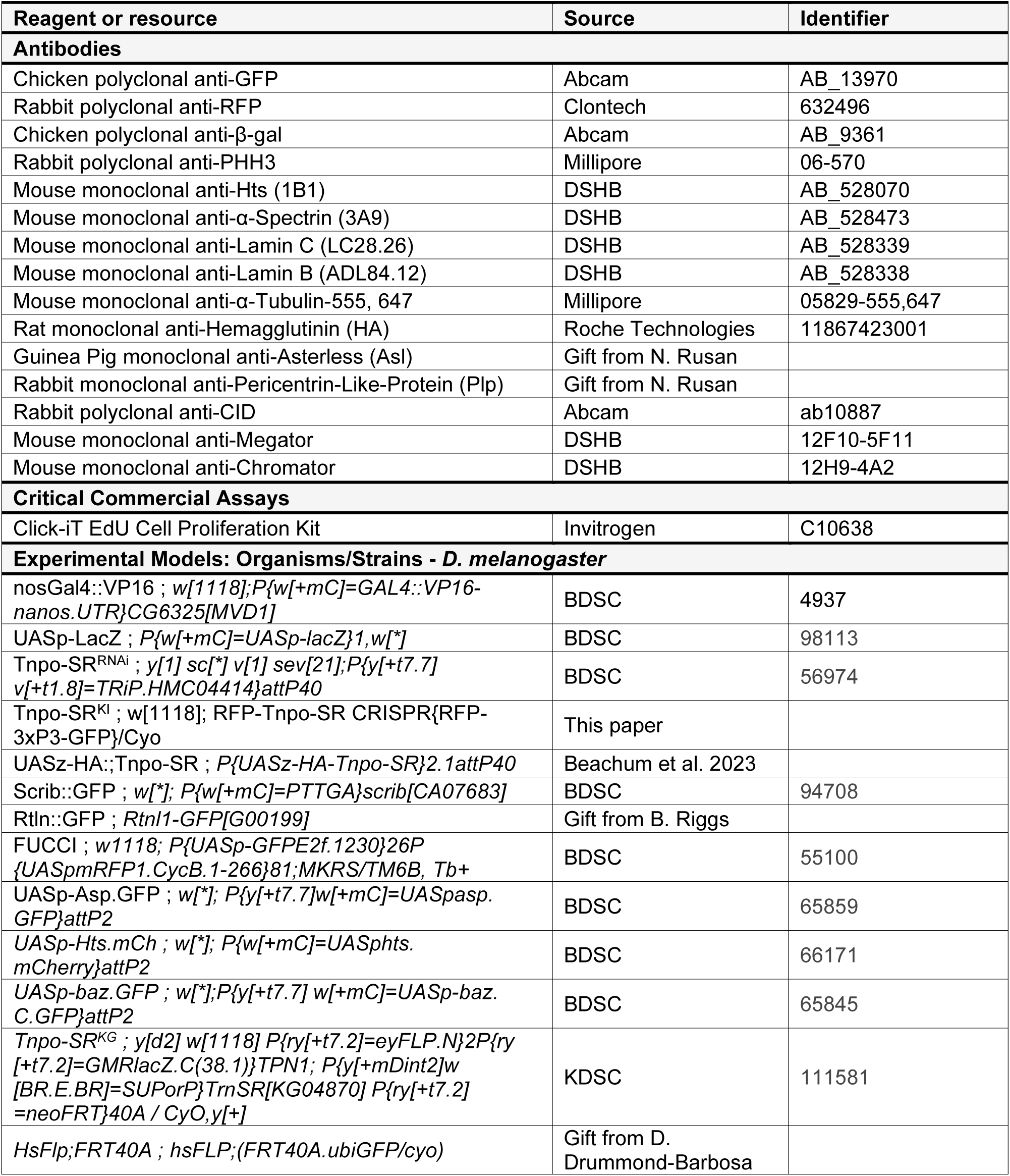

## SUPPLEMENTAL INFORMATION

**Figure S1:**
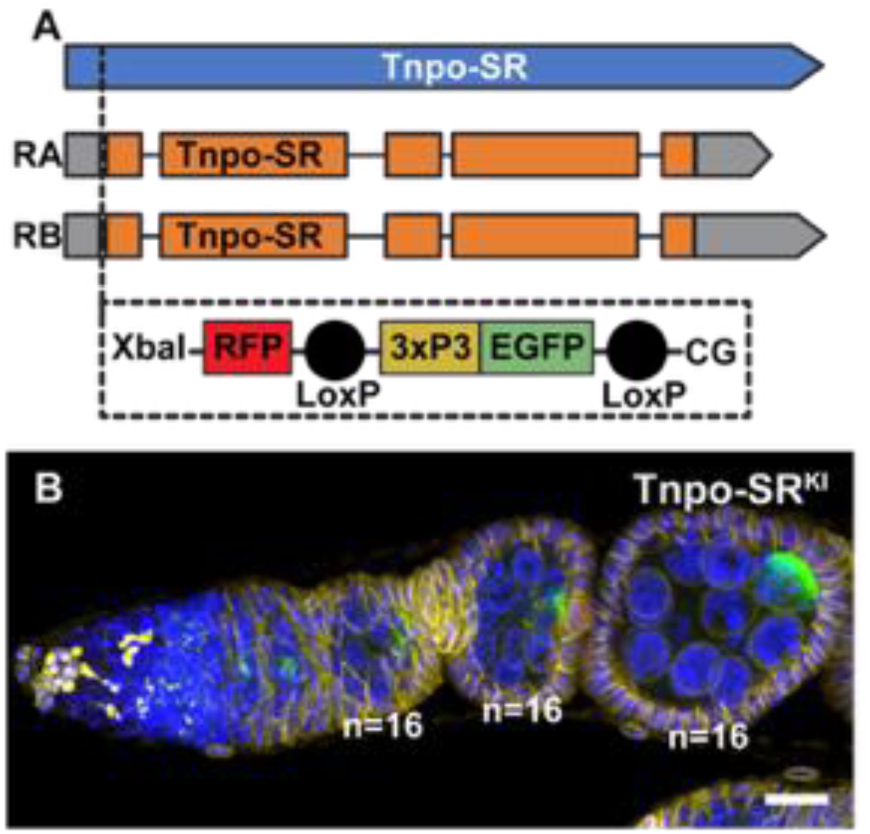
Generation and characterization of a *Tnpo-SR^KI^* allele. (A) Using CRISPR, an N-terminal RFP was inserted after the ATG in the endogenous *Tnpo-SR* locus. Cassette *RFP-3xP3-GFP* contains an Xbal site, RFP, and a floxed selection marker *3xP3-GFP* (LoxP – 3xPax3 promoter – EGFP – ⍺-Tub 3’ UTR – LoxP). *Tnpo-SR^KI^* with the selection marker is homozygous lethal and does not express RFP in ovarian cells. Flies carrying the knock-in construct after Cre-mediated excision of the selection marker (*mCherry::Tnpo-SR*) (Beachum et al., 2023) were also homozygous lethal and expressed RFP in ovarian cells. (B) Ovariole from *Tnpo-SR^KI^* heterozygous female immunostained for oocytes (Orb, green), the fusome (Hts, with LamC to mark cap cells, magenta), and DNA (DAPI, blue). Scale bar = 20µm.

**Figure S2:**
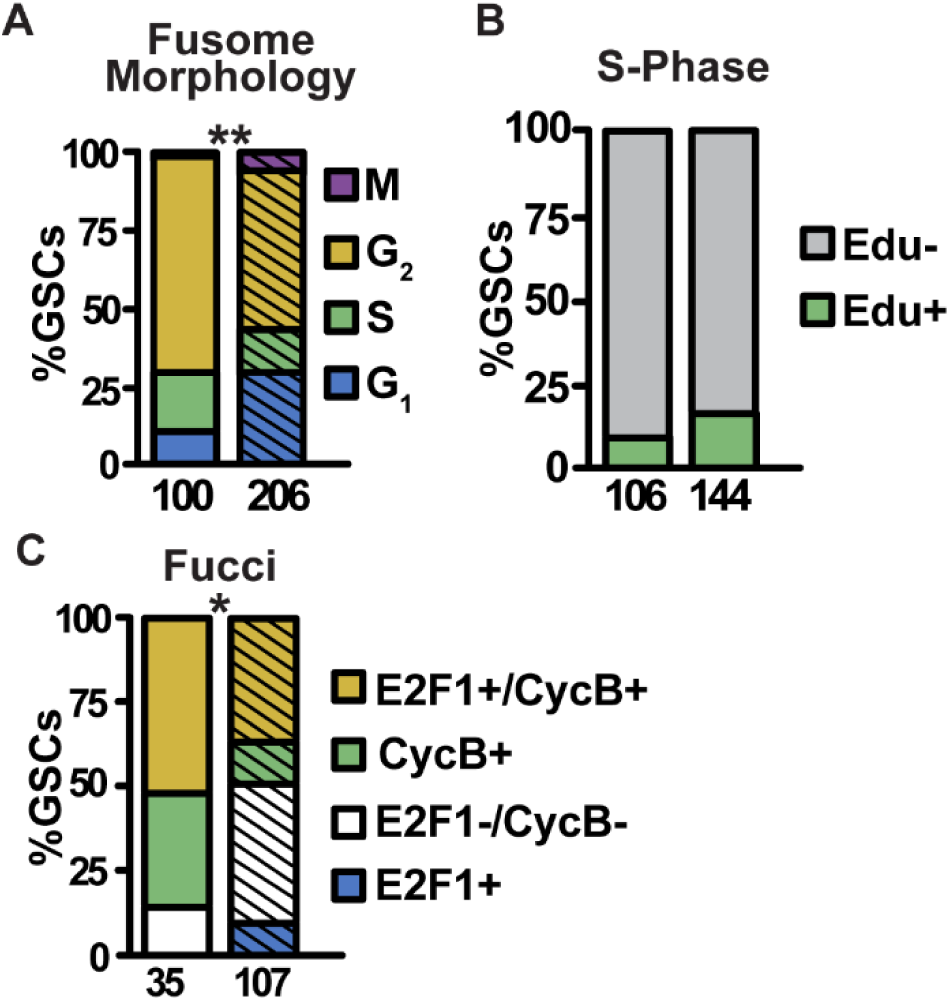
Depletion of *Tnpo-SR* delays exit from G1. Quantification of cell cycle progression via fusome morphology (A), incorporation of EdU in a one-hour pulse (B) or Fly-FUCCI fluorescent labeling (C) in control (solid) or *Tnpo-SR^RNAi^* (striped) GSCs. G_1_ (blue and white), S (green), G_2_ (yellow), M (purple). Mitotic GSCs were identified by chromosome morphology (labeled with DAPI). (B) Percentage of GSCs that are EdU positive (green) or negative (gray). Numbers below bars represent the number of GSCs analyzed. Significance determined by Fisher’s Exact test to compare the number of GSCs in G_1_ (plug morphology, A; E2F1+, C) or the number of EdU+ GSCs (B) to the total number of GSCs. \*\**p*<0.001, \**p*<0.01.

**Figure S3:**
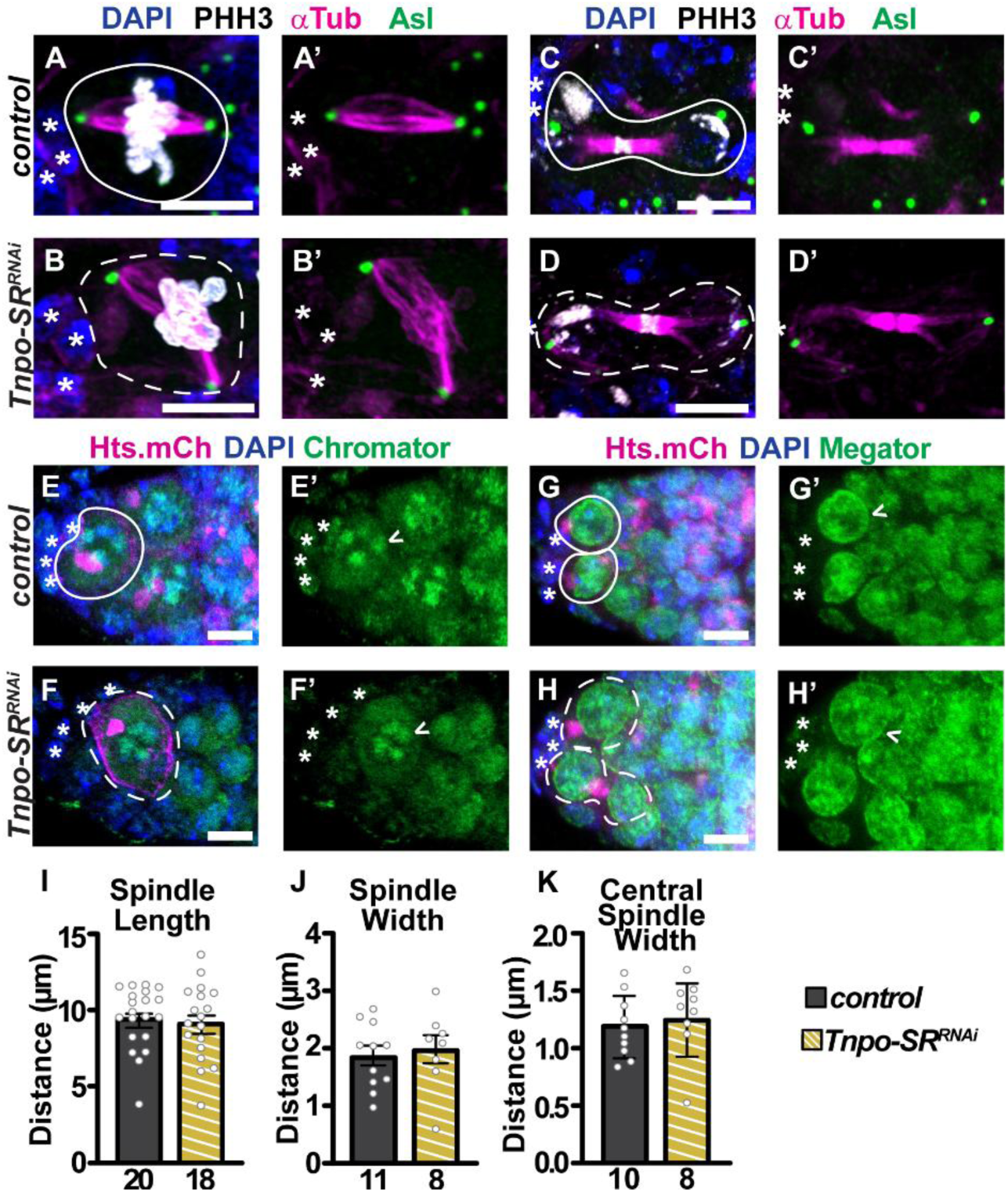
*Tnpo-SR* is dispensable for mitotic spindle and matrix formation. (A-H) control (A, C, E, G) and *Tnpo-SR^RNAi^* (B, D, F, H) germaria immunostained for a mitotic marker (PHH3, white; A-D), the fusome(Hts, white (C-D) or Hts.mCherry, magenta (E-H)), microtubules (⍺-Tubulin, magenta; A-D), centrosomes (Asl, green; A-D), spindle matrix proteins Chromator (green; E-F), Megator (green; G-H) and DNA (DAPI; A-H). GSCs outlined with white lines (solid-control, dashed-mutant). Asterisks denote cap cells. Scale bar = 5µm. (I-K) Quantification of GSC mitotic spindle length (I) and width (J) or central spindle width (K) in control (solid) or *Tnpo-SR^RNAi^* (striped, yellow) germaria. Numbers under bars represent the number of GSCs analyzed. Error bars represent s.e.m. Significance determined by Student’s two-tailed T-test.

**Figure S4:**
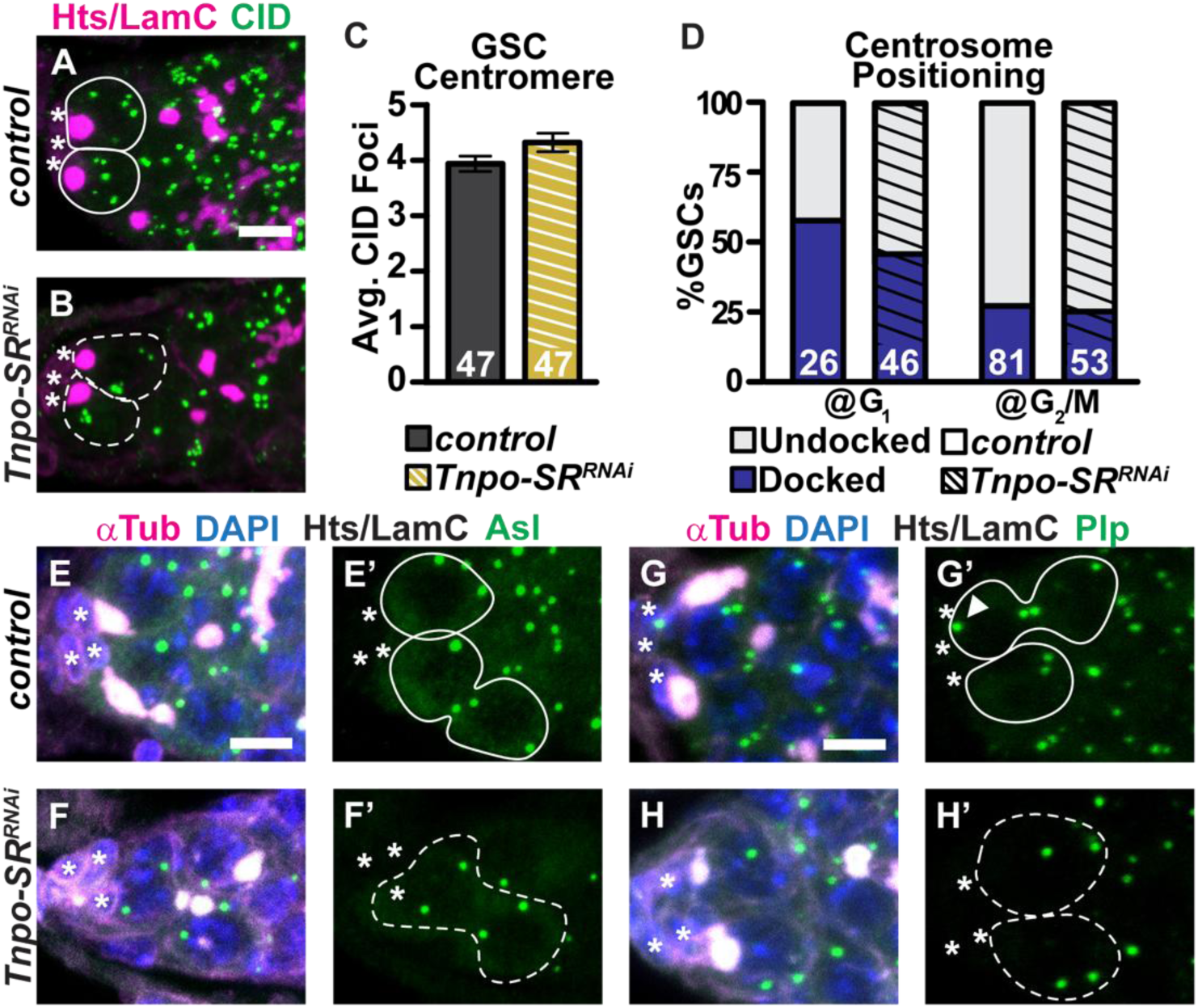
*Tnpo-SR* is not required to maintain centromere number or centrosome positioning. (A-B, E-H) Control (A, E, G) and *Tnpo-SR^RNAi^* (B, F, H) germaria immunostained for centromeres (CID, green; A-B), centrosomes (Asl, green; E-F), Plp, green; G-H) the fusome (Hts, magenta (A-B) or white (E-H)), microtubules (⍺-Tubulin, magenta; E-H) and DNA (DAPI, blue; E-H). (C) Average number of CID foci. (D) Percentage of GSCs with one centrosome docked at the somatic cap cells at G_1_ or G_2_/M phase of the cell cycle. Arrowheads denote docked centrosome. Asterisks denote cap cells. GSCs outline by white lines (solid-control, dashed-mutant). Numbers in bars represent the number of GSCs analyzed. Scale bar = 5µm. Error bars represent s.e.m. Significance determined by Student’s two-tailed T-test (C) or Fisher’s Exact test (D).

**Figure S5:**
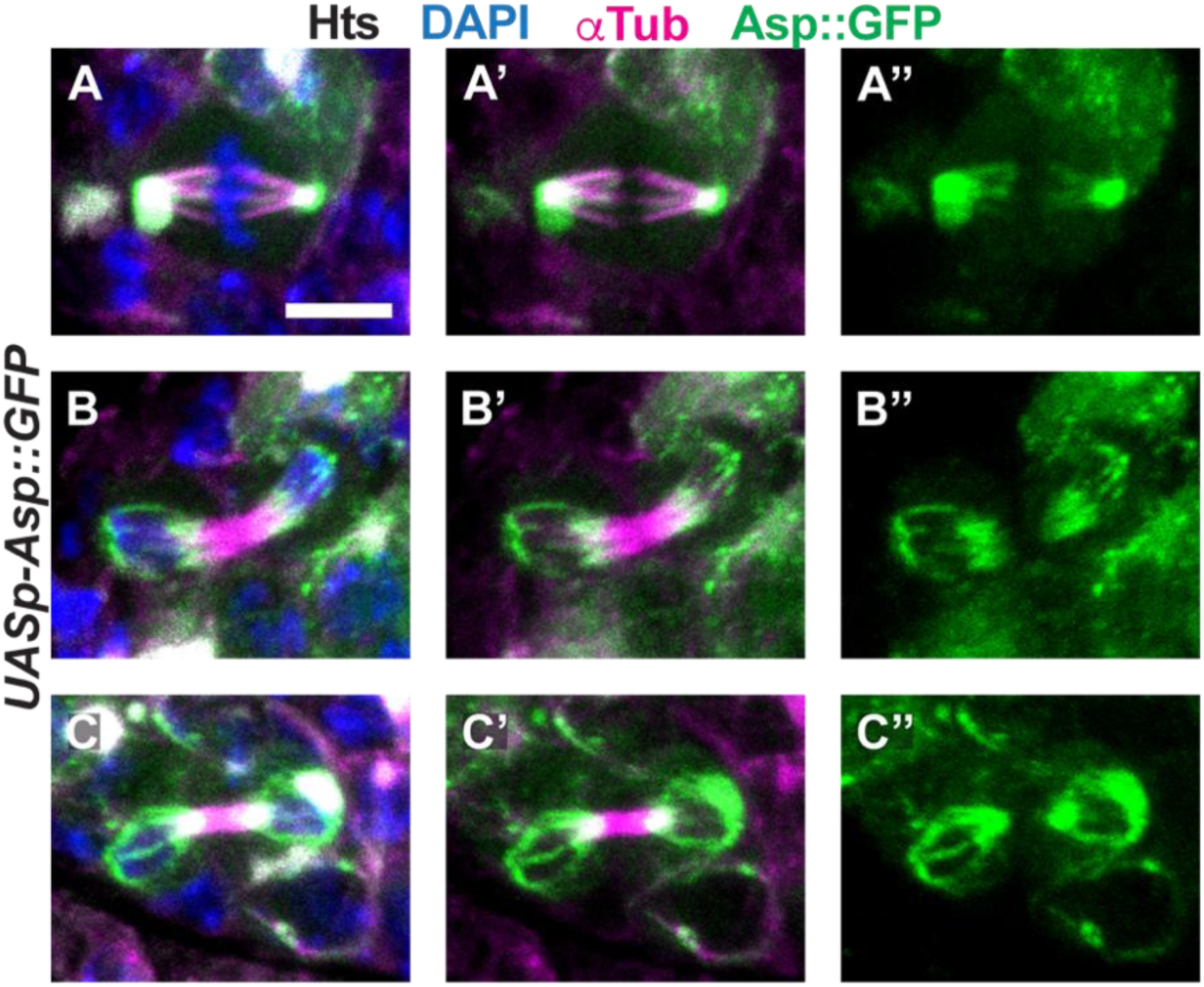
Asp localization at mitosis. (A-C) *UASp-Asp::GFP* germ cells immunostained for Asp (green), microtubules (⍺-Tubulin, magenta), the fusome (Hts, white), and DNA (DAPI, blue). Scale bar = 5µm.

## Notes

### Competing Interest Statement

The authors have declared no competing interest.

### Summary of Updates

Additional controls and analyses were added to this version of the manuscript, which altered some of the original conclusions (including the title). The text was reorganized and condensed, shifting some negative data to supplementary information.

